# Deacetylation of catalytic lysine in CDK1 is essential for Cyclin-B binding and cell cycle

**DOI:** 10.1101/420000

**Authors:** Shaunak Deota, Sivasudhan Rathnachalam, Kanojia Namrata, Mayank Boob, Amit Fulzele, S Radhika, Shubhra Ganguli, Chinthapalli Balaji, Stephanie Kaypee, Krishna Kant Vishwakarma, Tapas Kumar Kundu, Rashna Bhandari, Anne Gonzalez de Peredo, Mithilesh Mishra, Ravindra Venkatramani, Ullas Kolthur-Seetharam

## Abstract

Cyclin-dependent-kinases (CDKs) are essential for cell cycle progression. While dependence of CDK activity on Cyclin levels is established, molecular mechanisms that regulate their binding are less studied. Here, we show that CDKl:Cyclin-B interactions are regulated by acetylation, which was hitherto unknown. We demonstrate that cell cycle dependent acetylation of the evolutionarily conserved catalytic lysine in CDK1 or eliminating its charge state abrogates Cyclin-B binding. Opposing activities of SIRT1 and P300 regulate acetylation, which marks a reserved pool of CDK1. Our high resolution structural analyses into the formation of kinase competent CDK1: Cyclin-B complex have unveiled long-range effects of catalytic lysine in configuring the CDK1 interface for Cyclin-B binding. Cells expressing acetylation mimic mutant of Cdc2 in yeast are arrested in G2 and fail to divide. Thus, by illustrating cell cycle dependent deacetylation as a determinant of CDK1:Cyclin-B interaction, our results redefine the current model of CDK1 activation and cell cycle progression.

## Introduction

Cyclin-dependent kinases (CDKs) are evolutionarily conserved protein kinases, which are indispensible for eukaryotic cell division cycles, differentiation and transcription [1, 2], Among these, CDK1 (or yeast Cdc2 and Cdc28), CDK2, CDK4 and CDK6 orchestrate cell cycle progression. These constitutively expressed CDKs form cognate pairs with Cyclins, which have cell cycle phase dependent expression [1], Besides Cyclin binding, our current understanding of the temporal control of CDK activity is based on inhibitory and activatory phosphorylations, brought about by a regulatory loop involving other kinases and phosphatases [3]. Aberrant activation or inhibition of CDKs are known to cause cancer, developmental disorders and senescence [4].

Protein regions that are required for CDK-Cyclin interactions have been mapped [5-7], Cyclin binding induces the movement of PSTAIRE-helix in CDK and thus the formation of an active kinase conformation [6, 7], Interestingly, activatory phosphorylation, for example in CDK1, has been shown to stabilize the CDKl:Cyclin-B complex, although, this phosphorylation is itself dependent upon Cyclin-B binding to CDK1 [5, 8]. Given the essentiality of Cyclin binding for CDK activation and hence cell cycle progression, physiologically relevant mechanisms that determine complex formation remain to be addressed. Specifically, deterministic residues within CDKs that may regulate Cyclin binding/unbinding and therefore contribute to cell division are poorly understood. It is important to emphasize that Cyclin binding to CDKs have been generally thought to be default.

Almost all eukaryotic protein kinases (EPKs), including the CDKs, have a conserved catalytic triad consisting of N-lobe-β3 lysine, glutamate in C-helix and aspartate in DFG motif, which are essential for kinase activity [9, 10]. Several structure-function studies, mostly on cAMP dependent Protein Kinase A (PKA), have attempted to investigate the roles played by these catalytic triad residues in rendering the kinase active and in mediating phosphoryl-transfer. Uniquely in CDKs, binding of Cyclin induces a structural change to bring together these catalytic triad residues, which are otherwise far apart in apo-CDKs [6, 11]. Moreover, N-lobe-β3 containing the catalytic lysine is contiguous with the PSTAIRE-/C-helix, which not only harbors the catalytic glutamate but also forms the interface where Cyclin binds [6, 12]. Therefore, given that Cyclin binding brings a distinct mode of activation of CDKs, identification of regulatable residues or structural elements, which lead to the formation of the kinase competent complex will be crucial to our understanding of cell division cycles.

EPKs, including CDKs, have been reported to be acetylated [13-19]. It is important to note that in many of these cases, acetylation happens on the catalytic lysine. Substitution of the catalytic lysine to acetyl-mimic glutamine, for example in CDK2, CDK5 and CDK9, leads to a complete loss of kinase activity [14-16, 20]. This raises the possibility of acetylation as a key determinant of CDK activity, i.e. in addition to phosphorylation. Despite these studies, if and how acetylation and deacetylation dynamics regulate CDK activity has not been elucidated.

Here, we report that CDK1 is acetylated at lysine-33 by the acetyltransferase P300 and is deacetylated by NAD^+^-dependent deacetylase SIRT1. We show that while lysine-33 is dispensable for ATP binding, the conformation and interactions of ATP in the active site are altered upon its acetylation. Surprisingly, we find that perturbing the charge state of catalytic lysine impacts CDK1 and Cyclin-B complex formation. Moreover, we also show that CDK1 lysine-33 acetylation is dynamically regulated across cell-cycle stages and negatively correlates with CDKl:Cyclin-B complex formation. Substitution of the lysine to acetyl-mimic glutamine in Cdc2 failed to compliment different temperature-sensitive *cdc2* mutants in fission yeast Together, this study identifies deacetylation as a regulatory mechanism for activation of CDK1, which was hitherto unknown.

## Results

### SIRT1 and P300 regulate acetylation of the conserved catalytic lysine in CDK1

Given that CDKs are acetylated, if and how acetylation impinges on their structural and functional states is poorly understood. Specifically, the impact of lysine acetylation on CDK1 function, which in general cannot be compensated by any of the other CDKs [21], has not been addressed thus far. In this context, we probed CDK1 acetylation in mammalian cells. We found immunoprecipitated CDK1-HA to be acetylated (Fig 1A) consistent with our global LC-MS/MS analyses of the acetylated sub-proteome (unpublished results). This analysis revealed lysine-33 as the sole acetylation site in endogenous CDK1 (Fig IB), as reported earlier [13]. To further characterize the importance of this acetylation, we generated antibodies specific to the lysine-33 acetylated peptide sequence ^30^VAM**K**^Ac^KIRLESE^40^ in CDK1 (α-CDKl-K33Ac/α-K33Ac). This region is largely conserved across CDKs and specifically in CDK1 from yeast to humans (Figs S1A-S1C). Immunoaffinity purified α-K33Ac antibodies were able to detect endogenous CDK1 acetylation at lysine-33 (Fig S1E) and did not react with CDK1-K33R mutant (Fig S1D). Further, we could detect acetylated species of CDK1 using both pan-acetyl-lysine and α-K33Ac antibodies (Fig 1C). These clearly corroborated the LC-MS/MS results vis-à-vis acetylation of CDK1 at lysine-33. Our results suggest that endogenous CDK1 acetylation is low in abundance (Fig S1E) and it is important to note that even phosphorylation, the dominant and deterministic modification on CDK1, is sub-stoichiometric, across cell cycle phases [22, 23].

**Figure 1.**
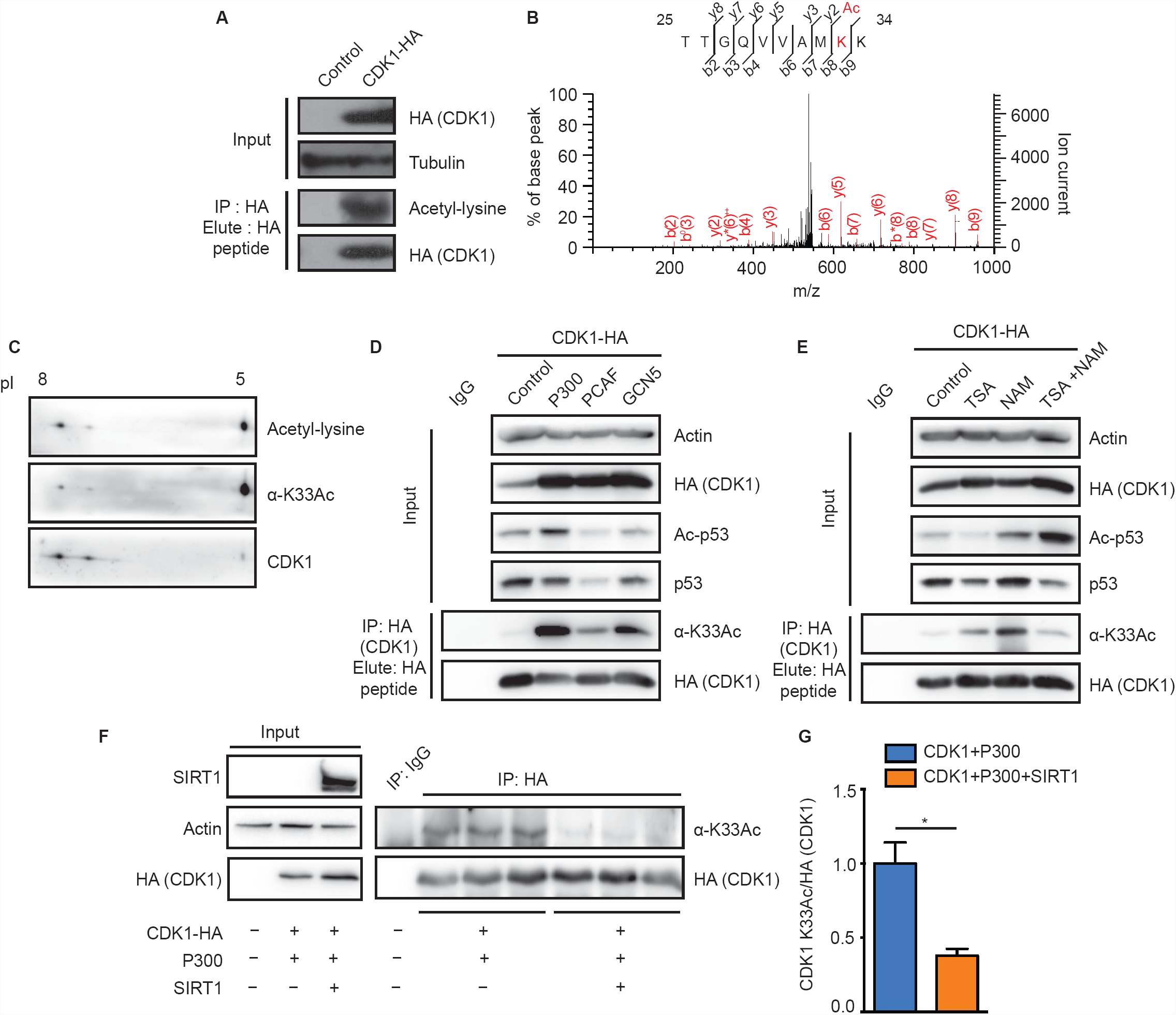
In-vivo CDK1 acetylation at lysine-33 is regulated by opposing actions of P300 and SIRT1. (A) Immunoprecipitated CDK1-HA was probed with a-pan-acetyl lysine antibody to reveal acetylation. (B) Identification of lysine-33 as the site of acetylation in CDK1 by LC-MS/MS analysis. (C) 2D-PAGE of immunoprecipitated endogenous CDK1 from HEK 293T cells and immunoblot with a-pan-acetyl-lysine or a-CDKl-K33Ac antibodies confirms acetylation. (D-E) Probing for acetylation of immunoprecipitated CDK1-HA from cells (D) cotransfected with P300, PCAF or GCN5, shows P300 as the major KAT and (E) treated with KDAC inhibitors indicates Nicotinamide (NAM) sensitive acetylation. (F-G) Probing for acetylation of immunoprecipitated CDK1-HA from cells cotransfected with P300 or P300/SIRT1 shows SIRT1 mediated attenuation of P300 dependent CDK1 lysine-33 acetylation. (F) Representative immunoblot and (G) Quantification of pixel intensities from (F). Error bars represent SEM for n=3 technical replicates from a representative experiment.

Protein acetylation is regulated by activities of lysine acetyltransferases (KATs) and lysine deacetylases (KDACs). Hence, to identify the acetyltransferase responsible for acetylating CDK1 at lysine-33, we expressed CDK1-HA along with KATs: P300, PCAF and GCN5. As can be seen from Fig ID, CDKl-acetylation was significantly enhanced in response to P300 expression when compared to PCAF and GCN5. Moreover, P300 was able to acetylate CDK1 *in-vitro* (Fig S1F). These results are consistent with earlier reports showing CDK1-P300 interaction and CDK1 mediated phosphorylation ofP300 [24],

To identify the KDAC, we treated cells with nicotinamide (NAM) and Trichostatin-A (TSA), which non-overlappingly inhibit Sirtuins (NAD^+^-dependent deacylases) and non-Sirtuin KDACs, respectively. We found that there was a robust increase in CDK1 acetylation following NAM treatment, clearly indicating an involvement of Sirtuins (Fig IE). Moreover, earlier reports have shown that CDK1 and SIRT1 interact with each other [25] (Fig S1G), although the biological relevance of this interaction has not been investigated. Given this, we indeed observe that SIRT1 overexpression led to a reduction in CDK1 acetylation (Figs IF and 1G). Taken together, these results clearly illustrate that P300 is the major KAT for CDK1 at lysine-33 and that SIRT1 is involved in its deacetylation.

### Acetylation of CDK1 at lysine-33 impairs its kinase activity

Evolutionarily conserved lysine-33 lies in the catalytic pocket of CDK1 where ATP binds, and analogous active site lysines in other CDKs and in fact all other kinases have been implicated in kinase activity [14-17, 26-33]. Hence, to check the importance of lysine-33 acetylation for CDK1 function, we employed K33R and K33Q mutants of CDK1, which have typically been used as deacetylation and acetylation mimics, respectively. *In-vitro* kinase assays of CDK1-WT and -K33Q, immunoprecipitated from G2/M synchronized cells coexpressing Cyclin-Bl (Fig S2B), showed that K33Q mutation completely abrogated the CDK1 activity (Figs 2A and 2B). Interestingly, CDK1-K33R also abolished kinase activity (Fig S2A) despite retaining the charge state, as with other kinases with similar arginine substitutions [17, 28-33]. It is important to note that although mutations of the catalytic lysine have been shown to abolish CDK activity, the underlying molecular mechanism is still unknown [14-16, 20, 27]. Notably, these results indicated that both the lysine side-chain and its charge state are essential determinants of kinase activity.

**Figure 2.**
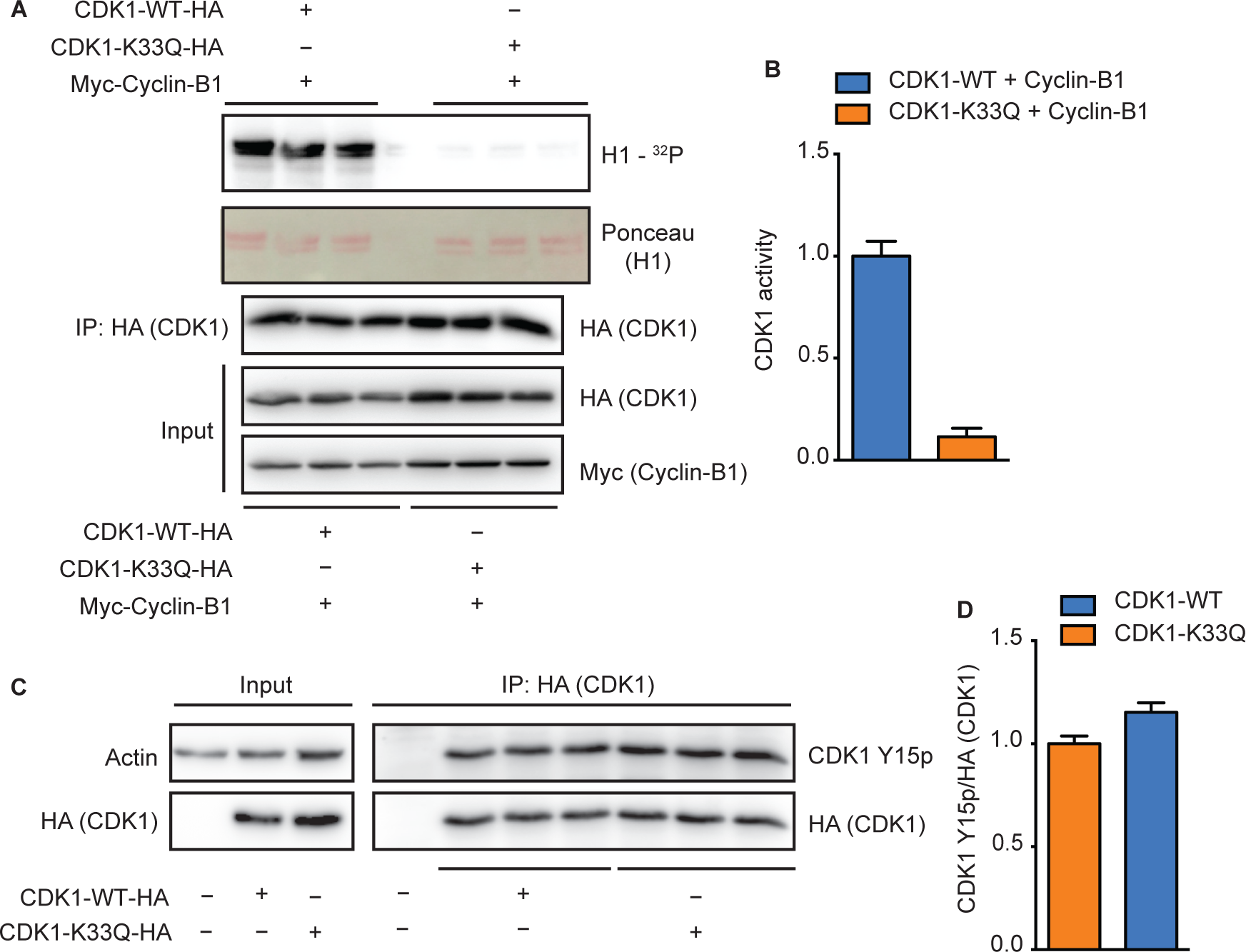
CDK1 acetylation at lysine-3 3 inhibits kinase activity. (A-B) HI phosphorylation by CDK1-K33Q expressed relative to the activity of CDK1-WT shows lysine-33 mutation abrogates kinase activity. Error bars represent SEM for n=3 technical replicates for two experimental repeats. (C-D) Immunoprecipitated CDK1-WT-HA or -K33Q-HA probed with a-phospho-tyrosine-15 CDK1 (CDK1 Y15p) antibodies reveals no change in inhibitory CDK1-Y15 phosphorylation. Error bars represent SEM for n=3 technical replicates from a representative experiment.

Mutations of tyrosine-15 and threonine-161 (inhibitory and activatory phosphorylation sites, respectively) or aspartate-146 (in DFG motif) in CDK1 affect cell cycle progression [26, 34-38]. Unlike these mutants, overexpression of CDK1-K33Q did not lead to a dominant negative phenotype (Figs S2C and S2D). Possibly owing to the kinase dead CDK1-K33Q mutant, we could not generate a cell culture system to assay for the acetyl-mimic mutant in a background wherein the endogenous wild-type CDK1 was absent (Data not shown). In this context, it is important to note that mutating an orthologous lysine residue to the acetyl-mimic glutamine (K40Q) in Cdc28 results in lethality [13]. Therefore, we resorted to using ectopically expressed CDK1-K33Q mutant to dissect out the molecular mechanism of loss of kinase activity.

Inhibitory phosphorylation at tyrosine-15 prevents precocious activation and is an indicator of inactive CDK1. Upon assaying for phospho-tyrosine-15, we did not find any significant difference between CDK1-WT and -K33Q (Figs 2C and 2D). This clearly rules out inhibitory phosphorylation as causal for inactivation of CDK1-K33Q mutant. Moreover, these motivated us to investigate a hitherto unknown mechanism of CDK1 activity regulation that is dependent on the catalytic lysine residue.

### Modeling and simulations of WT, K33Q and K33Ac CDKl:Cyclin-B:ATP complexes

Next, we investigated how lysine-33, its acetylation or the mutation to acetyl-mimic glutamine affected CDK1:Cyclin-B:ATP active complex, through modeling and computational analyses. To this end, we used the crystal structure of CDKl:Cyclin-B complex (PDB ID: Y472), reported recently [12], and docked an ATP molecule into the catalytic pocket We generated the wild type CDKl:Cyclin-B:ATP ternary complex model and validated it using the ATP bound CDK2:Cyclin-A crystal structure (Fig 3A), given their active site structural similarities. We also generated models for CDK1-K33Q and CDKl-K33Ac (lysine-33 replaced with acetyl-lysine) (Table SI). We would like to point out that the initial binding mode of ATP for all three models was the same. The atomistically detailed models were solvated, equilibrated at standard temperature (300 K) and pressure (1 bar), and subjected to 200 ns Classical Molecular Dynamics (CMD) simulations to relax the complexes to their native ternary form. Importantly, given that acetylated lysine is bulkier than lysine (Fig S3A), we employed three different equilibration protocols to study the impact of K33Ac on CDK1:Cyclin-B:ATP complex. Specifically, we obtained three distinct structurally relaxed conformations of the ternary complex after equilibration: CDKl-K33Ac-1 (ATP, E51 both constrained during equilibration), CDKl-K33Ac-2 (ATP unconstrained, E51 constrained during equilibration), and CDKl-K33Ac-3 (ATP constrained, E51 unconstrained during equilibration) (Figs 3B-3F and Figs S3B-S3F). We stress that no constraints were employed in the 200 ns production simulations. In the case of CDKl-K33Ac-1, but not in others, the acetyl-lysine residue flipped out of the active site pocket during production runs. In CDKl-K33Ac-2 and -K33Ac-3, the uncharged residue was observed to interact with the ATP nucleobase (Figs 3G-3Kand FigS3G-S3K).

**Figure 3.**
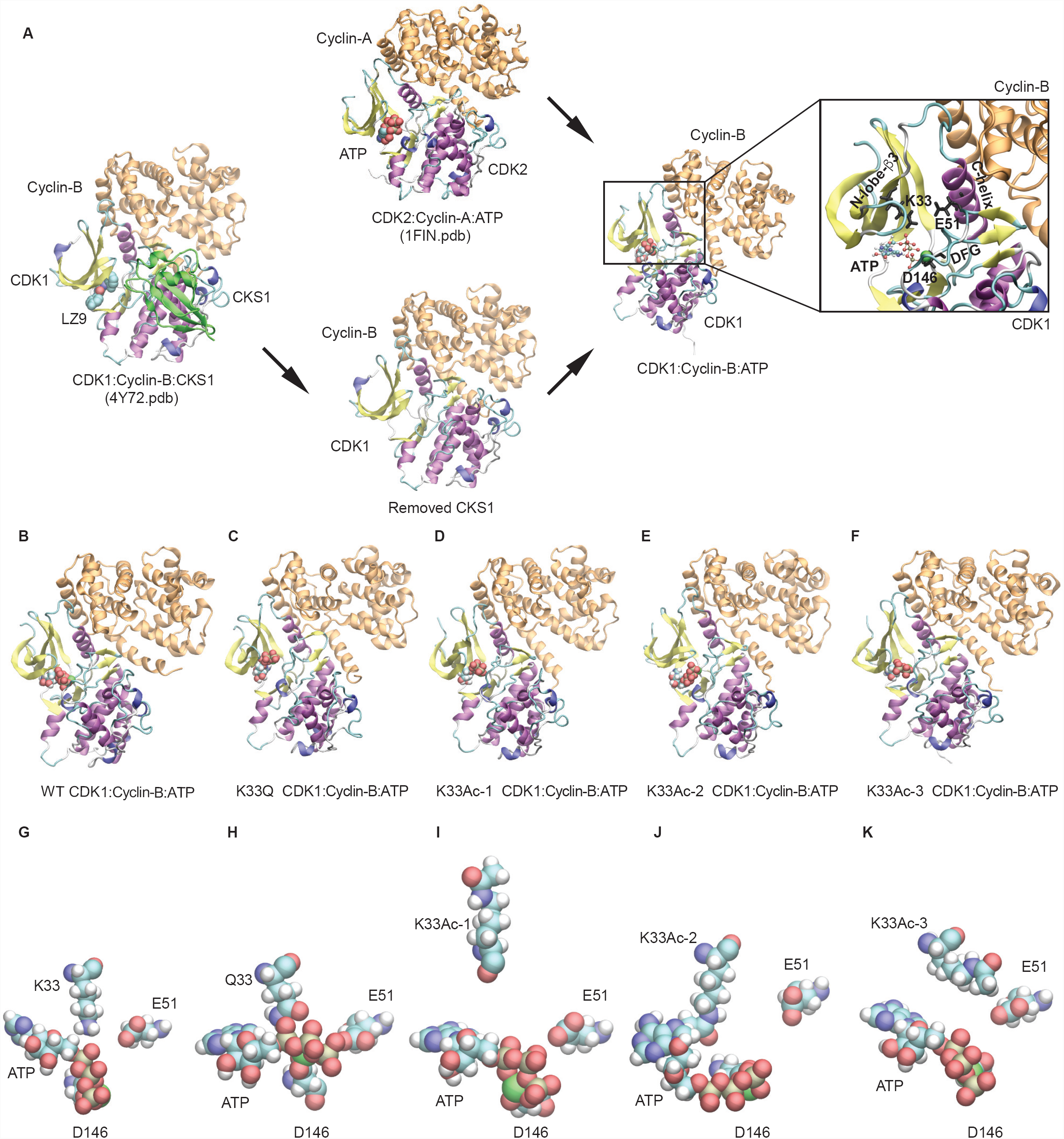
CMD simulations for WT, K33Q and K33Ac-l,-2,-3 CDKl:Cyclin-B:ATP ternary complexes. (A) Workflow depicting modeling of WT, K33Q and K33Ac-l,-2,-3 CDKl:Cyclin-B:ATP ternary complexes. (B-K) Structures for WT, K33Q and K33Ac-l,-2,-3 CDKl:Cyclin-B:ATP ternary complexes obtained from 200 ns NVT CMD simulations. (B-F) show CDK1 (multicolor) and Cyclin-B (orange) in secondary structure representation and ATP bound at the CDK1 active site is shown in atomic van der Waals spheres representation. (G-K) View of the active site showing key catalytic triad residues: lysine-33 (K33)/glutamine-33 (Q33)/acetyl-lysine (K33Ac), glutamate-51 (E51) and aspartate-146 (D146) interacting with the bound ATP and Mg^2+^ (green).

### Catalytic lysine in CDK1 plays distinct roles in stabilizing ATP and influencing interactions with Cyclin-B

We examined, *in-silico,* the effect of lysine-33 perturbations on ATP binding in terms of the following two measures: a) Solvent accessibility of ATP in the catalytic pocket relative to its accessibility in free solution (relative SASA), and b) Non-bonded (electrostatic and van der Waals) interaction energy of ATP with residues in the catalytic pocket (Fig 4B). As shown in Fig 4A, more than 60 % of the ATP surface area is buried in the active site in all five complexes (CDK1-WT, -K33Q, and the three -K33Ac models). Although in the CDKl-K33Ac-2 model, we observed transient fluctuations in the relative SASA for ATP during the 200 ns MD trajectory, the averaged values over the entire timescale show that ATP is buried in the active site as in the other models. Nevertheless, we found that with the loss of electrostatic interactions of ATP tail with lysine-33, as in CDK1-K33Q and -K33Ac complexes, the conformation adopted by ATP was different when compared to CDK1-WT (Fig 4D). We further assessed the non - bonded interaction energy (E_NB_) of ATP with residues of the active site pocket within a 3-7 Å radius around the ATP. In the wild type complex, for residues within 3 Å, E_NB_ was about -400 kcal/mol and become slightly more favorable (to around -450 kcal/mol) when residues up to 5 Å were included (Fig 4B and Fig S3L). In CDK1-K33Q and -K33Ac complexes, the ATP E_NB_ values were comparable and were less favorable (by about 100-200 kcal/mol), vis-à-vis CDK1-WT for interactions with residues within a 3-7 Å radius. Moreover, our analyses clearly suggested that interactions of ATP with residues beyond 5 Å appear to be insignificant, leading to negligible changes in E_NB_ for both CDK1-WT and mutant complexes.

**Figure 4.**
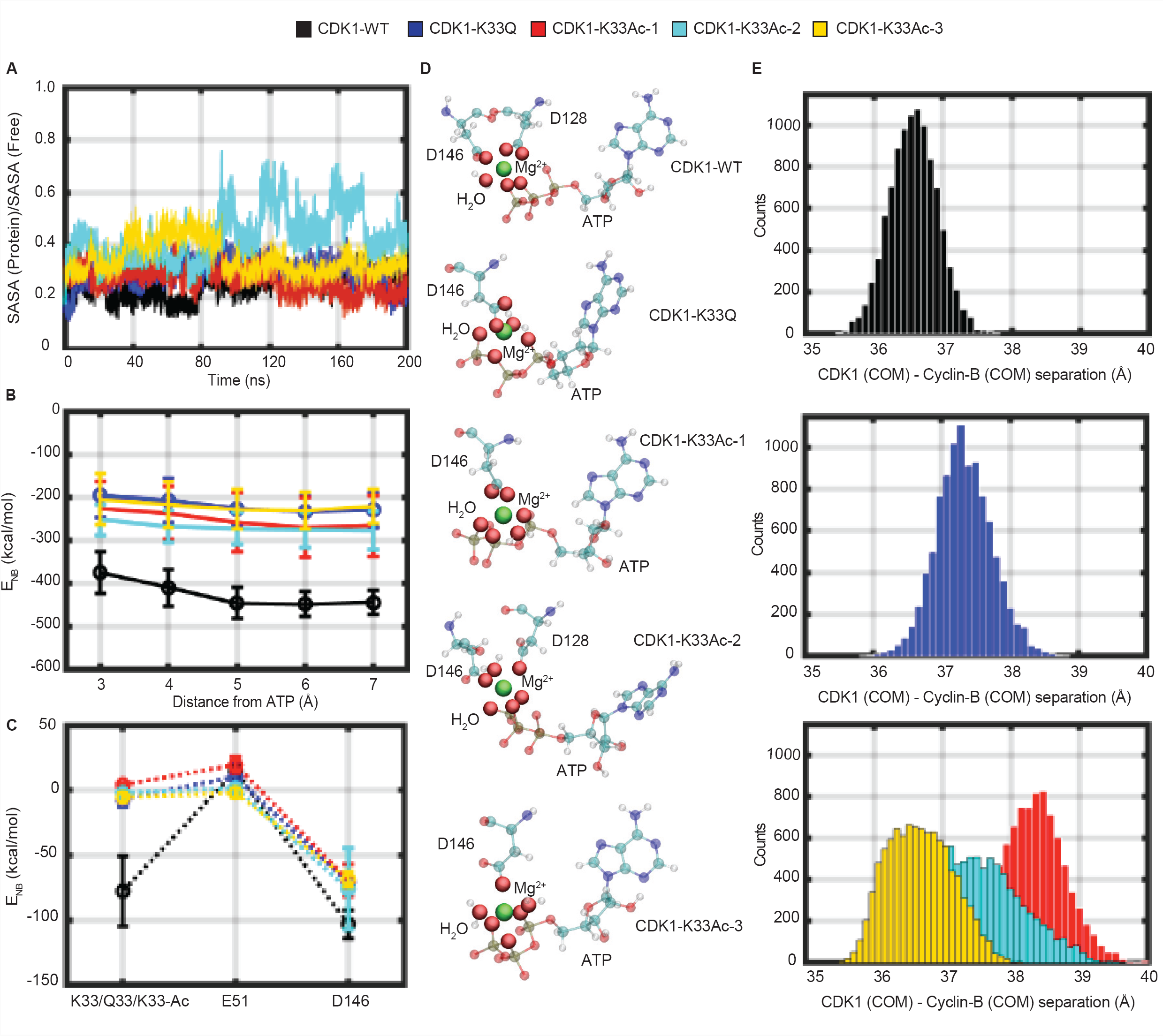
Loss of CDK1 lysine-33 charge state impinges on active site ATP conformation and weakens CDKl:Cyclin-B complexation. **(A)** Fractional Solvent Accessible Surface Area (SASA) for ATP (ratio of values in ternary complex and free solution) obtained during 200 ns NVT CMD trajectories for WT, K33Q and K33Ac CDK1:Cyclin-B:ATP systems are compared. **(B)** Comparison of non-bonded interaction energies (E_NB_) of ATP with surrounding active site residues within a 3-7 Å radius in WT, K33Q and K33Ac CDKl:Cyclin-B:ATP systems. Error bars represent SD. **(C)** A comparison of non-bonded interaction energy (E_NB_) of ATP with catalytic triad residues, residue-33 (K33, Q33, K33Ac), aspartate-146 (D146), and glutamate-51 (E51) in WT, K33Q and K33Ac CDKl:Cyclin-B:ATP systems highlights significant contributions from K33, which are reduced upon mutation. Error bars represent SD. **(D)** ATP orientations and Mg^2+^ (green) coordination spheres (including solvation) in WT, K33Q and K33Ac CDKl:Cyclin-B:ATP complexes. The ATP active site conformation and Mg^2+^ coordination are altered upon mutating lysine-33. **(E)** Histograms of Centre of Mass (COM) distances between CDK1 and Cyclin-B reveal greater separations and broader distributions of distances in K33Q and K33Ac CDK1:Cyclin-B:ATP complexes relative to WT.

An analysis of ATP interactions with the core catalytic triad (residue-33, aspartate-146, and glutamate-51) showed that lysine-33 was one of the major contributors in terms of binding energetics (Fig 4C). However, it should be noted that despite the loss of favorable non-bonded interactions, as in the case of CDK1-K33Q and -K33Ac, the overall interactions (E_NB_ < -200 kcal/mol) of ATP with the residues of active site pocket remain highly favorable (Fig 4B). In summary, ATP appears to be stably bound to the same extent, albeit with different binding modes in CDK1-WT, -K33Q and -K33Ac complexes. In particular, we find that the Mg^2+^coordination sphere, which includes oxygens of ATP, acidic residues at the active site and water molecules to be different for CDK1-WT, -K33Q and -K33Ac complexes (Fig 4D). Based on these results on lysine-33 perturbations in CDK1, and the previous report of lysine-72 mutations in PKA [33], we propose that the catalytic lysine may be dispensable for stable ATP binding in EPKs. Nevertheless, our analyses raise the possibility of lysine-33 to play a key role in phosphoryl-transfer reaction, which needs to be addressed in the future.

Interactions between the catalytic residues lysine and glutamate on N-lobe-β3 and C-helix, respectively, have been proposed to be important for kinase activity in almost all EPKs [10, 39]. Specifically, in a complex kinase system like the CDKs, Cyclin binding induces a salt-bridge interaction between the catalytic lysine and PSTAIRE-/C-helix glutamate [7, 12, 40]. Importantly for CDKs, the impact of the loss of lysine charge state on salt-bridge with glutamate and PSTAIRE-/C-helix interactions has not been investigated. In this regard, for CDK1-WT we found that lysine-33 interacted with both glutamate-51 and ATP during the time course of the simulations, showing more prominent interactions with the latter (Figs S3M-S30). The change in the lysine-33 charge state caused a small rotation of the PSTAIRE-helix axis by ∼5 degrees in the CDK1: Cyclin-B interfacial plane (Figs S4A and S4B), and also changed the position and conformation of the T-loop (activation segment) in the CDK1 catalytic domain (Figs S4C and S4D). However, we did not find any movement of either the PSTAIRE-/C-helix or glutamate-51 towards the ATP because of the loss of lysine-33 charge state (Figs S3P-S3R), unlike what was recently reported for PKA [33]. This indicates that the catalytic lysine, while being indispensable, plays differential roles in rendering kinases active across EPKs.

Mutational analyses and structural evidences have indicated that PSTAIRE-/C-helix in CDK1 is involved in binding to Cyclin-B, similar to CDK2:Cyclin-A [6, 7, 12, 41]. However, the forces/interactions that stabilize the CDK: Cyclin complex remain unclear. In our attempts to examine the consequences of loss of lysine-33 charge state, we discovered a change in the distribution of center-of-mass (COM) distances between CDK1 and Cyclin-B proteins in the CDK1-K33Q and -K33Ac complexes relative to the CDK1-WT. As can be clearly seen in Fig 4E, the distribution of CDK1: Cyclin-B COM separations for the mutant lysine-33 complexes (CDK1-K33Q, -K33Ac-1 and -K33Ac-2) distinctly shift to larger values relative to CDK1-WT. For CDKl-K33Ac-3, while there is no significant shift in peak COM values, the distribution broadens significantly, indicating higher flexibility in the binding geometry of the two proteins. In fact, the widths of distribution of the inter-protein separations along our MD trajectories are the smallest for CDK1-WT. These observations suggest that the perturbations of lysine-33 could be involved in remodeling CDKl:Cyclin-B interface, which was unknown thus far.

To summarize, our *in-silico* structural analyses using multiple CDKl:Cyclin-B:ATP ternary complexes led us to hypothesize that: a) While the charge state of lysine-33 in the catalytic pocket was necessary for orienting ATP, it may be dispensable for its binding and b) Perturbing lysine-33 could influence CDK1 interactions with Cyclin-B. We therefore set out to validate these hypotheses using biochemical approaches.

### Catalytic lysine-33 is dispensable for ATP binding

To check if lysine-33 is required for ATP binding, we used the following approaches: a) binding of 5’-(4-Fluoro-sulfonyl-benzoyl)-adenosine (FSBA), an analog of ATP and b) ATP-agarose in which ATP is conjugated to the beads in different orientations. Consistent with previous reports from CDK9 [14] and Cdc2 [42], CDK1-K33Q and -K33R mutants had reduced FSBA binding (Fig 5A). FSB A covalently binds to the active site lysine with high affinity [43]. Moreover unlike FSBA, ATP is non-covalently coordinated in the catalytic pocket. Thus FSBA binding that is equivalently affected in CDK1-K33Q and -K33R does not reveal the importance of lysine in ATP binding.

**Figure 5.**
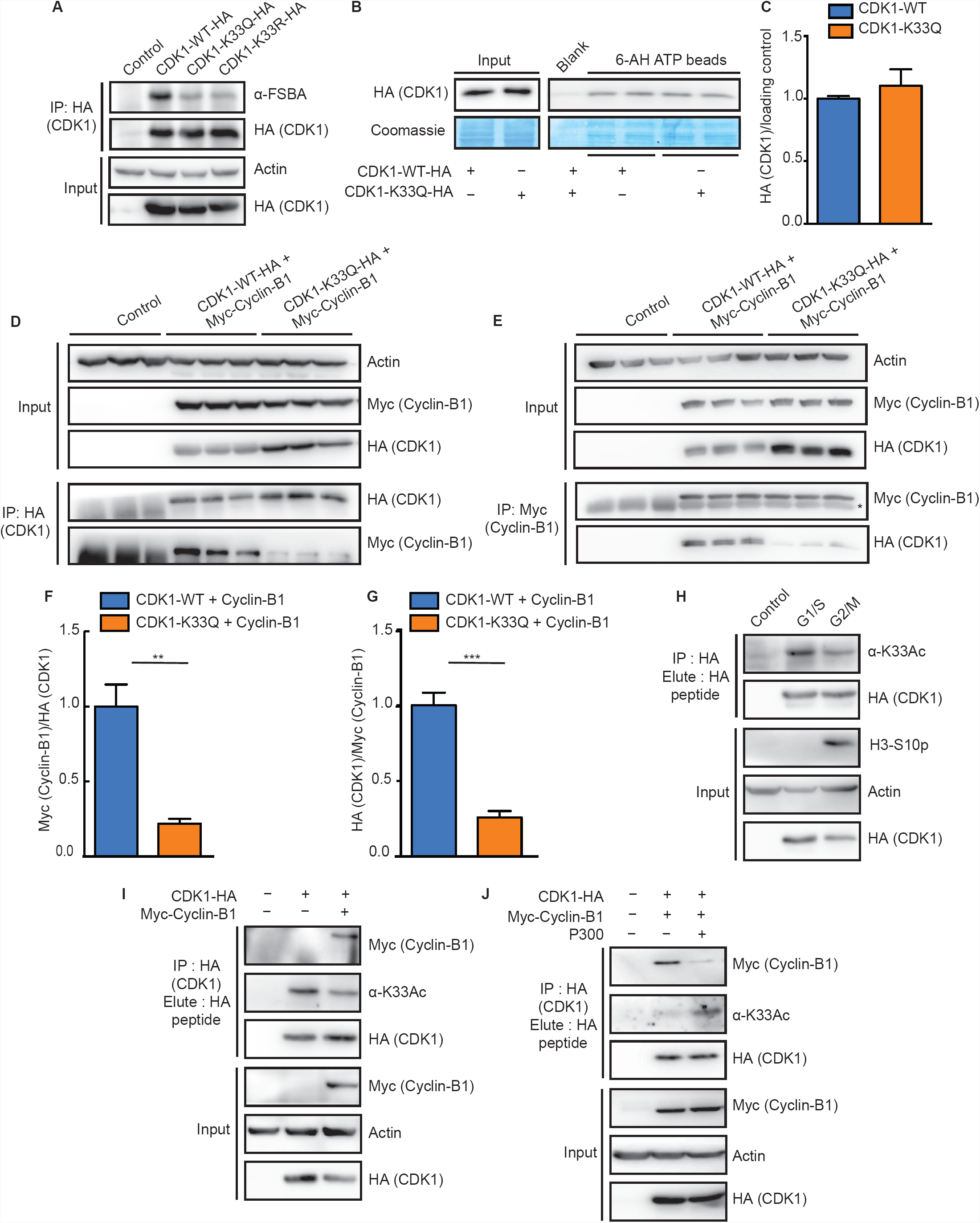
CDK1 acetylation at lysine-33 does not affect ATP binding but reduces Cyclin-Bl interaction. (A) Probing FSBA labeled CDK1-WT/-K33Q/-K33R-HA immunoprecipitates with α-FSBA antibodies revealed reduced binding upon K33 mutation. (B) Probing for CDK1-WT-HA and -K33Q-HA binding (pull-down) by ATP conjugated agarose beads (6-AH beads) shows no change in steady-state binding. Coomassie staining of total bound proteins was used for normalization. (C) Quantification of ATP binding in (B). Error bars represent SEM for n=5 technical replicates from two experimental repeats. (D-G) Reversible co-immunoprecipitations of CDK1-WT-HA or -K33Q-HA and Myc-Cyclin-Bl shows reduced CDKl:Cyclin-Bl binding upon K33Q mutation. Immunoprecipitation of complexes with (D) α-HA and (E) α-Myc antibodies. (F) and (G) Quantifications of results presented in (D) and (E), respectively. *marks the IgG heavy chain. Error bars represent SEM for n=3 technical replicates from a representative experiment. (H) Probing for acetylation in immunoprecipitated CDK1-HA shows reduced CDK1 lysine-33 acetylation in G2/M synchronized cells as compared to Gl/S phase. Phospho-H3 S10 (H3-S10p) was used as a marker for G2/M population. (I) Immunoprecipitated CDK1-HA shows negative correlation between CDK1 lysine-33 acetylation and CDK1:Cyclin-Bl complexation on probing with α-K33Ac and α-Myc antibodies. (J) Probing immunoprecipitated CDK1-HA with α-K33Ac and α-Myc antibodies from cells transfected with Myc-Cyclin-Bl or Myc-Cyclin-Bl/P300 shows enhancing lysine-33 acetylation reduces Cyclin-Bl binding.

We found that CDK1-WT bound 6-AH ATP-beads better than the others and were used for further analyses (Fig S5A). Next, we assessed the relative binding efficiency of CDK1-WT, -K33Q and -K33R to the ATP conjugated beads. In contrast to our results with FSBA, we found that steady-state ATP binding is not affected by these mutations in CDK1 (Figs 5B and 5C and Figs S5B and S5C). Nevertheless, in binding competition assays using ATP in solution, we found small differences between CDK1-WT, -K33Q and -K33R (Figs S5D and S5E). These results clearly indicate that the catalytic pocket lysine or its charge state are dispensable for steady-state ATP binding and importantly corroborate our predictions based on the *in-silico* analyses.

### Non-default binding of Cyclin-B to CDK1 is determined by the charge state of catalytic lysine

Having ruled out all known possible mechanisms that led to CDK1 inactivation due to the alterations of catalytic lysine and motivated by our *in-silico* results, we wondered if Cyclin-B binding itself is affected. To test this, we probed for the interaction of CDK1-WT, -K33Q and -K33R with Cyclin-Bl by reversible co-immunoprecipitations. Remarkably, we found that CDK1-K33Q, which takes away the charge and is typically used as an acetylation mimic, reduced the interaction with Cyclin-Bl (Figs 5D-5G). On the contrary, CDK1-K33R, which retains the charge state and mimics constitutively deacetylated state, remained bound to Cyclin-Bl (Figs S5F and S5G).

Given that we found endogenous CDK1 lysine-33 to be acetylated, which would naturally mask its charge state, we wanted to assess if acetylation is a determinant of Cyclin-B binding in a cell cycle dependent manner. Despite our best attempts, we could not map differential acetylation of endogenous CDK1 during different phases of cell cycle using α-K33Ac antibodies (Data not shown). As is known for most posttranslational modifications, including key regulatory phosphorylations [22, 23], stoichiometry of protein acetylation has been shown to be relatively low [44, 45]. Therefore, our inability to quantitatively estimate cell cycle dependent changes in acetylation could be due to low abundance of acetylated CDK1 combined with the low sensitivity of α-K33Ac antibodies. Nevertheless, using over-expressed CDK1, we investigated if the acetylation and deacetylation potential was indeed regulated in a cell cycle dependent manner. As shown clearly in Fig 5H and Fig S5H, we found hypoacetylated CDK1 in G2/M synchronized cells when compared to cells arrested in Gl/S. Importantly, overexpression of Cyclin-Bl, which increases G2/M population, also resulted in reduced CDK1 lysine-33 acetylation (Fig 5I). This was consistent with our hypothesis of deacetylation of catalytic lysine as a determinant of Cyclin-B binding. We further assayed for CDKl:Cyclin-Bl complexes under conditions, which led to increased CDK1 lysine-33 acetylation, by immunoprecipitation. As anticipated, we found that hyperacetylated CDK1 showed significantly less Cyclin-Bl association (Fig 5J). We consistently failed to express and purify soluble CDK1 and Cyclin-Bl, independently (Data not shown), given that recombinant CDK1: Cyclin-B are only obtained by co-expressing them together. Due to this limitation, we could not carry out *in-vitro* binding studies to assess strength of interaction of CDK1-K33Q or -K33Ac with Cyclin-B, which are complexation defective. Together, we have found CDKl:Cyclin-B interactions to be regulated, which was hitherto unknown. Importantly, our findings establish that the positive charge of catalytic lysine-33 in CDK1 is essential for Cyclin-B binding and its loss either by acetylation or a mutation to glutamine abrogates the interaction.

### Electrostatic tethering of catalytic lysine-33 dictates surface interactions between CDK1 and Cyclin-B

Next, we wanted to investigate the mechanistic underpinnings of lysine-33 dependent association of CDK1 and Cyclin-B proteins. Examining the structure of ternary WT CDKl:Cyclin-B:ATP complex revealed that residues 40-60 of CDK1 interface directly with Cyclin-B. Specifically, in addition to the PSTAIRE-helix (residues 45-58), which has been previously described [6,12], we found that the random coil segment (residues 40-44) in CDK1 also interacts with Cyclin-B, as implicated in CDK2:Cyclin-A [6]. Importantly, the random coil segment is contiguous with the N-lobe-β3 (residue 28-36), containing lysine-33. It should be noted that combination mutants of residues in each of these segments in CDK1 have been shown to reduce Cyclin-B binding [5]. However, the forces that orient these segments and help in stabilizing interactions between the proteins are still unknown.

These observations prompted us to investigate if the local electrostatic interactions of catalytic lysine-33 impinge on interface interactions of CDK1 with Cyclin-B. Specifically, we carried out a statistical analysis of salt-bridge interactions in CDK1-WT, -K33Q and -K33Ac ternary complexes (Table S2) during the initial (first 1 ns) stages of our 200 ns MD trajectory (Fig 6C). We chose the 1 ns period of the trajectory since destabilization of the CDK1:Cyclin-B interface, as assessed by inter-protein COM separations, occurred early in our MD production runs in CDK1-K33Q and -K33Ac systems. Our analysis clearly shows that the number of interfacial salt-bridges is halved in the case of CDK1-K33Q and -K33Ac complexes, as compared to the CDK1-WT, during the initial phase of our MD simulations (Fig 6A), presaging the increase in inter-protein separations (Fig 4E). Further, we quantified the distributions of E_NB_ of interfacial salt-bridge forming residues of CDK1 and Cyclin-B (Table S2). Interestingly, we found the E_NB_ values to be significantly lower (by ∼150-200 kcal/mol) and also highly variable when the charge state of lysine-33 is lost (Fig 6B). Moreover, a more detailed analyses revealed that interfacial salt-bridge interactions involving both, acidic (D/E) and basic (K/R) residues of CDK1 (Fig S6A), as well as side chain-backbone salt-bridges (Fig S6B) were reduced in K33Q and K33Ac CDKl:Cyclin-B:ATP complexes.

**Figure 6.**
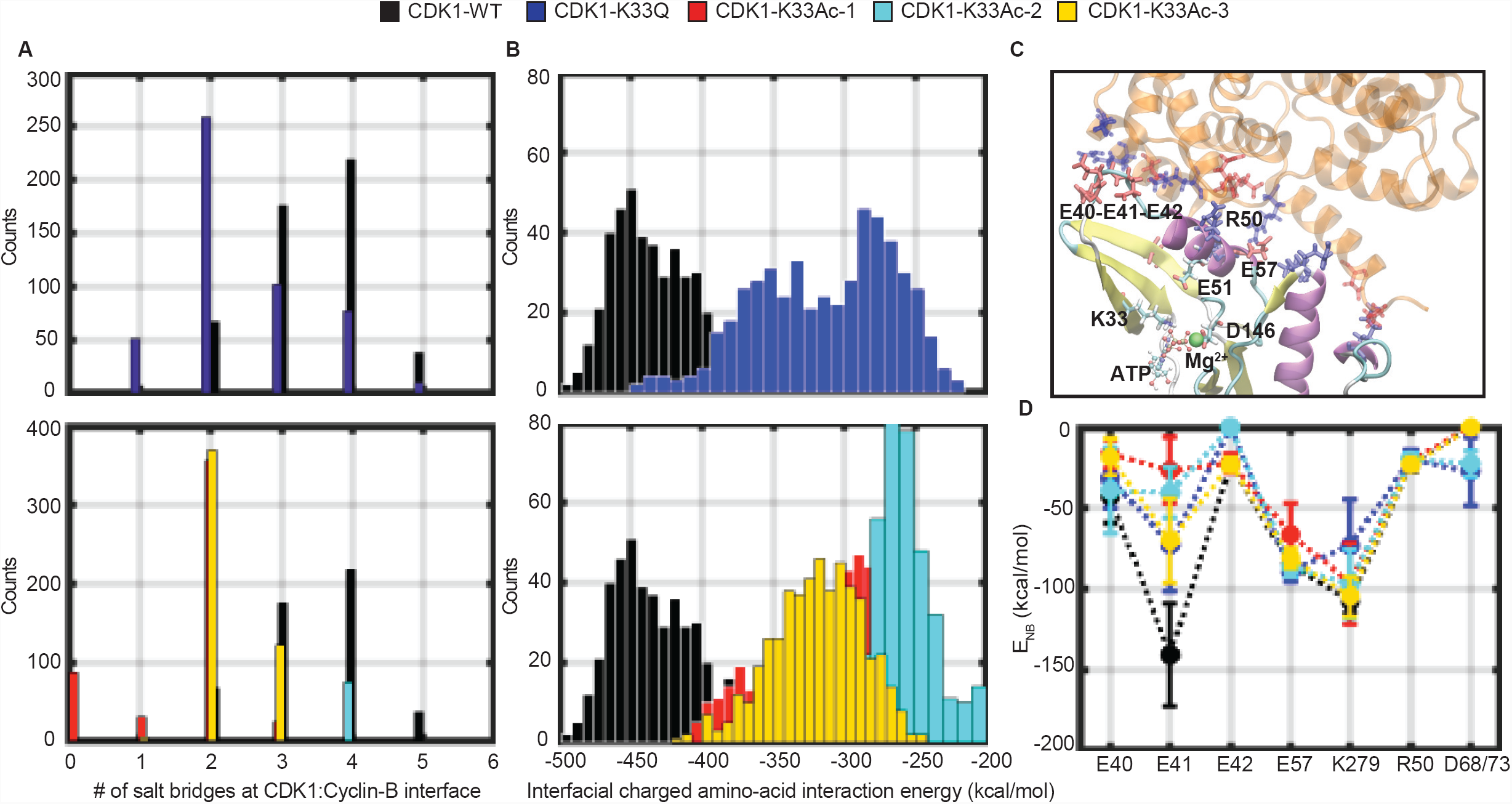
Loss of CDK1 lysine-33 charge state affects surface electrostatic interactions with Cyclin-B. (A) Reduction in the number of interfacial salt bridges between CDK1 and Cyclin-B in CDKl:Cyclin-B:ATP K33Q (top panel) and K33Ac systems (bottom panel) relative to WT. (B) Reduction in non-bonded interaction energies between acidic and basic residues at CDKl:Cyclin-B interface in CDKl:Cyclin-B:ATP K33Q (top panel) and K33Ac systems (bottom panel) relative to WT. (C) View of the CDK1: Cyclin-B interface and its connectivity to the active site with catalytic triad residues (K33, D146, E51) and ATP. Key acidic and basic residues at the CDK1: Cyclin-B interface are also shown. In this representation, we show only residues 30-148 and 250-280 in CDK1. (D) Comparison of non-bonded interaction energies (E_NB_) of key acidic and basic residues of CDK1 with salt bridge partner residues of Cyclin-B shows maximum contribution by E41, which is dependent on K33 charge state. Error bars represent SD. The analysis is for data from the beginning of first 1 ns of NVT production runs for WT and mutant systems before CDKl-Cyclin COM-COM separations increase for the latter.

Our comprehensive analysis further demonstrated that, upon perturbing lysine-33, the largest changes in E_NB_ were observed for residue glutamate-41 (by ~100 kcal/mol) at the CDKl:Cyclin-B interface (Fig 6D). Interestingly, we found negligible perturbations in salt-bridge interactions formed by acidic and basic residues in the PSTAIRE-helix (arginine-50 and glutamate-57). Specifically, our results illustrate that the effects mediated by catalytic lysine-33 on glutamate-41 in the N-lobe-β3/C-helix linker loop of CDK1 may play a deterministic role in enabling CDK1: Cyclin-B interactions. These findings are also illustrated in the form of movies (Movies S1-S5).

### Cdc2/CDK1 lysine-33 mutation blocks cell-cycle progression in *S. pombe*

To better understand physiological consequence of lysine-33 acetylation for CDK1 function, we turned to fission yeast, *Schizosaccharomyces pombe,* which shares the cell cycle regulatory mechanisms with higher eukaiyotes including humans [46, 47] and is one of the best studied system for mitotic control. Indeed the first human CDK gene was cloned based on its ability to compliment a temperature sensitive mutation in *cdc2* gene, *cdc2*-33 [46]. Unlike mammalian cells, the fission yeast cell cycle is controlled by a single CDK, Cdc2, which drives both Gl/S and G2/M transitions, making it a simple model to study cell cycle progression [48-50]. Moreover a single CDK/Cyclin pair (Cdc2p/Cdcl3p) is sufficient to drive progression through the different phases of cell cycle [51].

Wild type fission yeast cells grow to a length of about 14 microns before dividing. Temperature-sensitive (ts) *cdc2* mutants that undergo cell-cycle arrest at G2 and are unable to divide or form colonies, continue to grow when incubated at the restrictive temperature of 36°C producing abnormally elongated cells with a single nucleus (Fig 7 A). Two independent ts alleles of *cdc2, cdc2-33* (A177T) and *cdc2-L7* (P208S), [48, 52] arrest as elongated mononucleate cells when incubated at restrictive temperature of 36°C (Figs 7B and 7C and Figs S7A-S7D), and fail to form viable colonies (Figs 7D and S7E). Expressing wild type *cdc2* gene under moderate strength thiamine repressible promoter (pnmt41) rescues the cell cycle arrest in these mutants. These cells grow and divide at wild type length distribution and are viable at the restrictive temperature of 36°C (Figs 7B-7D and Figs S7A-S7E). Unlike the cells rescued with Cdc2-WT, inducible expression of both Cdc2-K33Q and Cdc2-K33R failed to compliment the two *cdc2* ts mutants. These cells were elongated, arrested in G2 and also failed to form viable colonies (Figs 7B-7D and Figs S7A-S7E). Although, an earlier study in *S. cerevisiae* had also shown that Cdc28-K40Q and Cdc28-K40R failed to complement *cdc28* null cells vis-à-vis viability [13], the effect on cell cycle was not addressed. Importantly, the findings described in the current study are consistent with our hypothesis about the significance of the catalytic lysine-33, which is evolutionarily conserved, in rendering CDK1/Cdc2 kinase competent and hence its role in cell cycle progression.

**Figure 7.**
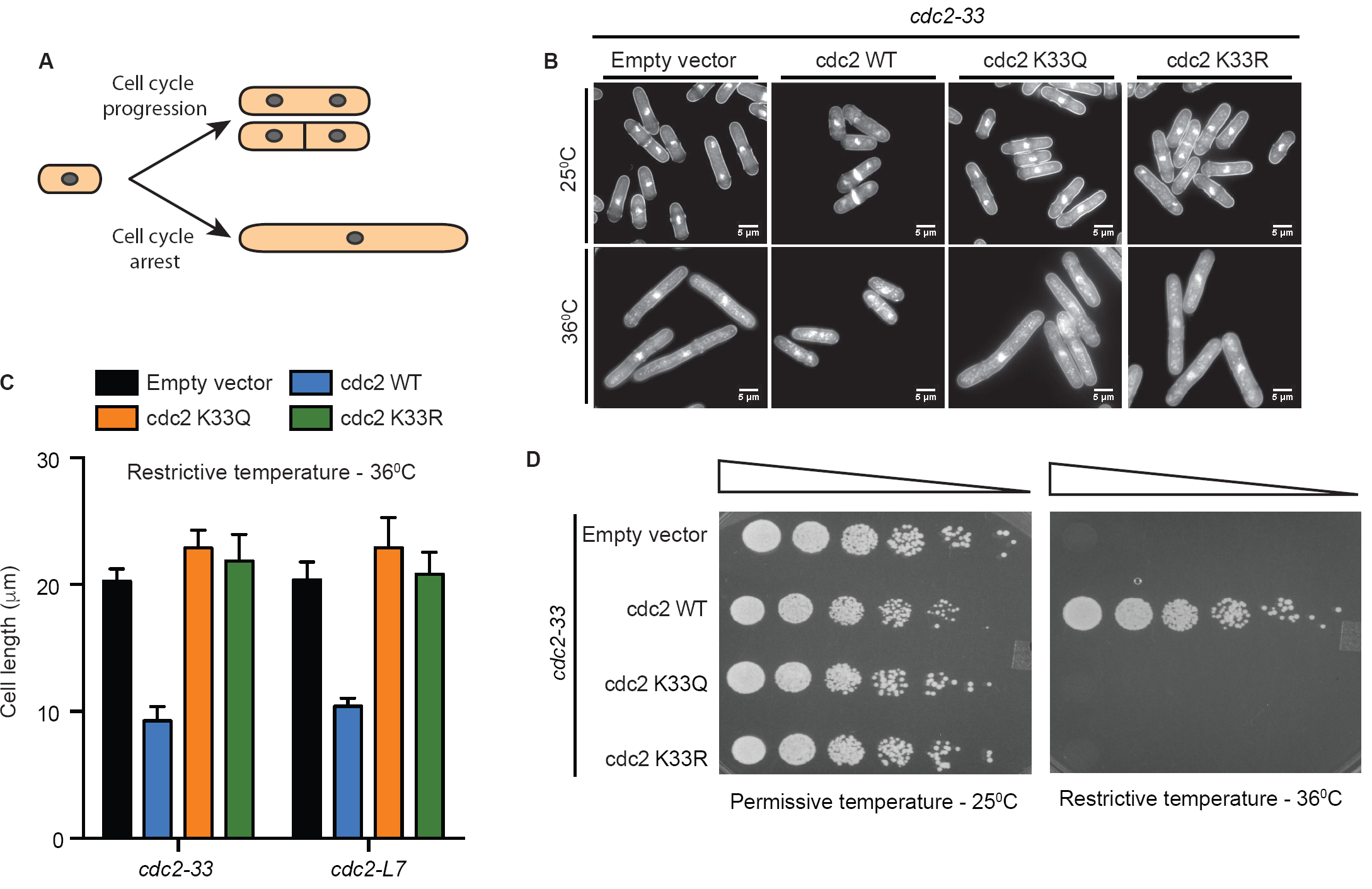
Cdc2/CDK1 lysine-33 mutation blocks cell-cycle progression in 5. *pombe* and fails to compliment *cdc2* ts mutants. (A) Schematic representation of cell length observed in fission yeast temperature sensitive *cdc2* mutants at permissive temperature (normal cell cycle progression) and restrictive temperature (cell cycle arrest). (B) Representative images of DAPI and aniline blue co-stained cells of fixed samples after 16 h of induction, before (top panel) and after (bottom panel) shift at restrictive temperature for 4 h. Cells fixed after 4 h incubation at 36°C. (C) Quantification of cell length measurements observed in *cdc2-33 and cdc2-L7* strains over-expressing indicated plasmids at restrictive temperature for 4 h. Measurement indicates mean cell length and SEM for three independent experiments where n=500 for each sample. (D) Plate growth assay after 16 h of induction. Samples were 1:4 serially diluted and plates were incubated at indicated temperature for 4-5 days.

## Discussion

In this study, we establish the molecular mechanism of acetylation-dependent inhibition of CDK1 and its role in cell cycle progression. Protein acetylation has been largely shown to occur on lysine residues in proteins [53-55] and our results indicate CDK1 to be mono-acetylated. We identified P300 and SIRT1 as key enzymes that regulate CDK1 acetylation. Importantly, our results illustrate that CDK1 is hypoacetylated in G2/M synchronized cells as compared to those in Gl/S. Earlier reports have shown that both SIRT1 and P300 interact with CDK1:Cyclin-B and the latter interaction led to phosphorylation-dependent degradation of P300 [24], Our results now provide physiological significance for interaction between these key molecular components in regulating CDK1 acetylation and thus cell division cycles.

Capturing the importance of CDK1 deacetylation, mutating lysine-33 to glutamine that mimics a constitutively acetylated state, resulted in a kinase dead CDK1. Although, CDK1-K33R retained the charge state, it was inactive and mechanistically it was distinct from CDK1-K33Q (see below). These suggested that the charge state and/or shape complementarity of the catalytic lysine are necessary for kinase activity. Notably, overexpression of CDK1-K33Q in the background of endogenous CDK1 did not lead to a dominant negative effect, unlike the phosphorylation defective mutants [36]. Given that CDK1 knockdown has been shown to have variable effects on mammalian cells [56] and to eliminate possible compensatory role of CDK2 [57], we addressed the importance of catalytic lysine acetylation using *S. pombe.* As stated earlier and well established in the field, Cdc2 and CDK1 are highly conserved both at structural and functional levels [46]. Highlighting the physiological relevance of catalytic lysine and its charge state, we found that mutating Cdc2 lysine-33 led to cell cycle arrest and reduced viability in *S. pombe.*

Structural studies and evidence based on ATP analogues have led to the assumption that the catalytic lysine is critical for both ATP binding and phosphoryl-transfer reaction [14-17], In this context, our high resolution computational analyses and ATP-binding assays show that ATP is stably bound to CDKl:Cyclin-B even when lysine-33 is mutated to glutamine or replaced with acetyl-lysine. Given that most of the current understanding of kinase catalytic triads has come from PKA, including a very recent study [33], we compared the results on PKA with those on our CDKl:Cyclin-B complex. Unlike in PKA, our results show that perturbing lysine-33 does not alter the position of PSTAIRE-/C-helix vis-à-vis ATP. Nevertheless, we do show that in CDK1-K33Q and -K33Ac, ATP adopts distinct conformations in the active site pocket and the non-bonded interaction energies become less favorable relative to CDK1-WT. These insights distinguish the role of catalytic lysine in different kinase systems, which was previously unknown.

Oscillation of Cyclin levels during different cell cycle phases have been thought to be sufficient to form cognate complexes with CDKs to drive cell cycle progression [3]. Although, previous structural and mutational analyses of CDK:Cyclin complexes have elucidated the domains that mediate interactions [5,-7, 41], deterministic molecular mechanisms that enable complexation remained unclear. In this context, we have discovered a key role for catalytic lysine-33 in CDK1 in determining Cyclin-B binding using both computational and biochemical approaches, with an evident physiological effect on cell cycle progression. Specifically, mutating lysine-33 to acetyl mimic glutamine (K33Q) abrogated Cyclin-Bl binding to CDK1. Interestingly, this interaction was intact in CDK1-K33R, indicating that the charge state is necessary for enabling Cyclin-Bl binding. Further, this also shows that absence of kinase activity and cell cycle arrest in K-R mutants is independent of control via Cyclin-B binding. This could be due to abrogated phosphoryl-transfer reaction, as has been reported in other kinase systems including PKA [33]. We emphasize that none of the previous studies have dissected out the link between the catalytic lysine in CDKs and the interface interactions with Cyclins. Our exciting results highlight that intra-molecular electrostatic tethering of lysine-33 predisposes CDK1 to bind to Cyclin-B, which eventually leads to the formation of active kinase conformation. Thus, we establish that CDK: Cyclin interaction is regulated and determined by the catalytic lysine, contrary to our current understanding of this complexation to be default.

In conclusion, our results suggest that acetylation of CDK1, which is sub-stoichiometric and prevents Cyclin-B binding, marks a reserve pool of CDK1. Our findings highlight that deacetylation, which naturally unmasks the charge state of lysine-33, is a key determinant of CDKl:Cyclin-B binding. Considering that other CDKs are also acetylated at similar active site lysine, acetylation dependent regulation might be a universal phenomenon of CDKs and could be true for other EPKs as well. Importantly, our study highlights how local perturbations involving a key catalytic residue have long-range effects on protein-protein interactions.

## Materials and methods

### Cell Culture and transfection

HEK293T, HeLa-TetOn and Mouse embryonic fibroblast (MEF) cells were grown in DMEM high glucose medium (Sigma-D777) supplemented with 10% fetal bovine serum (Gibco) or 10% newborn calf serum (Gibco) and 1% antibiotic-antimycotic solution (Gibco), and maintained under standard 5% CO_2_ conditions. Cells were transfected using Lipofectamine 2000 (Life technologies) or Fugene 6 (Roche) as per manufacturer’s instructions. For inducible expression, 2 μg/ml doxycycline (Sigma) was added to the medium as indicated. For inhibiting HDACs or Sirtuins, cells were treated with 400 nM Trichostatin A (TSA) or 5 mM Nicotinamide (NAM), respectively, for 16 h. Sf21 *(,Spodoptera frugiperda)* insect cells (Thermo Fisher) were grown in Graces’ Media containing antibiotic solution (Sigma) and 10% fetal bovine serum (Gibco) at 25°C.

### Yeast and bacterial culture

*Schizosaccharomyces pombe* strains were transformed with *pREP41* plasmid encoding wild type or mutant *cdc2* gene using fast lithium acetate transformation method [58]. All strains were cultured in YE or EMM (MP Biomedicals) media with necessary supplements. 10 μM thiamine in EMM was used to suppress the *nmt41* promoter. For induction, cells initially cultured in EMM media containing thiamine were washed and subsequently cultured in EMM media without thiamine at 25°C. After 16 h of induction, the cultures were shifted to restrictive temperature of 36°C for 4 h *E.coli DH5a cells* were cultured in LB media supplemented with 100 μg/ml ampicillin for amplification of plasmids. 2% agar was used for solid media.

### Cell synchronization

Cells plated at 50% confluency were treated with 150 μM mimosine (Sigma) for 16 h to synchronize the cells in Gl/S phase. To obtain cells synchronized in different phases, mimosine treated cells were washed twice with PBS and released into fresh medium. Cells collected at 2-3 h, 5-6 h and 7-8 h post release corresponded to S-phase, G2-phase and G2/M-phase respectively, as assessed by flow cytometery and phospho-H3 (S10) levels. To synchronize cells in G2 phase, 9 μM R0-3306 (Sigma) was used for 16 h and for G2/M synchronization, cells were treated with Nocodazole (100 ng/mL) for 8 h after mimosine release.

### Flow cytometry

Cells were collected by trypsinization, washed with PBS and fixed using chilled 70% ethanol, overnight at -20°C. The fixed cells were washed with thrice with PBS, treated with RNase A (Sigma) and stained using propidium iodide (Sigma) for 30 minutes at 37°C. Cell-cycle profiles were acquired on a FACS Fortessa instrument (BD Biosciences) and analyzed using FACS Diva Software 6.0 (BD Biosciences).

### Cell lysate preparation

Cells were harvested and washed twice in ice cold PBS. Cells were lysed either in TNN (50 mM Tris-HCl pH 7.5, 150 mM NaCl, 0.9% NP 40, 1 mM PMSF, PhosSTOP, Protease inhibitor cocktail) or RIPA buffer (50 mM Tris pH 8.0, 150 mM NaCl, 0.1% SDS, 0.5% Sodium deoxycholate, 1% Triton X 100, 0.1% SDS, 1 mM PMSF, PhosSTOP, Protease inhibitor cocktail) for 15 minutes on ice. When probing for acetylation, buffers were supplemented with 4 μM TSA and 5 mM Nicotinamide to inhibit the activity of deacetylases. The lysate was cleared by centrifugation at 12,000 rpm for 15 minutes at 4°C and the supernatant collected as cell lysate. The protein concentration was measured using BCA kit (Sigma) as per manufacturer’s protocol.

### Immunoprecipitation and HA peptide elution

Cell lysates were pre-cleared with either Protein-A/-G sepharose beads (Sigma) for 1 h at 4°C on rotor. Equal amount of protein was taken and the cell lysates were incubated overnight with the indicated antibody. Equal volume of Protein-A/-G beads were added and incubated for 2 h at 4°C on rotor to pull down the immune complexes and subsequently washed thrice with the TNN lysis buffer for 10 minutes at 4°C on rotor. The washed beads were then boiled in SDS gel loading buffer to elute the bound proteins. Alternatively, for eluting CDK1-HA immunecomplexes, beads were incubated in HA peptide elution buffer (50 mM Tris pH 8.0, 150 mM NaCl, 100 μg/ml HA peptide) for 2 h at 27°C and with 1000 rpm mixing. The beads were pelleted and supernatant was boiled in SDS gel loading buffer.

### FSBA labeling

CDK1-HA immunoprecipitates on beads were treated with 50 μM FSBA (Sigma) in 0.1% PBS-Triton-X-100 for 15 minutes at 30°C with constant shaking, washed four times with 0.5% PBS-Triton-X-100 for 5 minutes at4°C on rotor and boiled in SDS gel loading buffer to elute the bound proteins. Western blotting was performed using 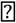-FSBA antibodies and normalized to the amount of immunoprecipitated CDK1.

### CDK1 kinase activity assay

Myc-tagged Cyclin-Bl was co-expressed with either CDK1-WT/-K33Q/-K33R-HA in HEK293T cells. 24 hr post-transfection, the cells were treated with Nocodazole (100 ng/mL) for 12 hr and harvested in TNN lysis buffer. The lysate was incubated with α-HA antibody overnight at 4°C, followed by Protein-A sepharose beads for 2 hr at 4°C. The beads were washed thrice with TNN buffer for 10 min each at 4°C, twice with kinase buffer (50 mM Tris-Cl pH 7.5, 15 mM MgCl_2_) for 10 min each at 4°C, and divided in two parts, one for western blotting to estimate the amount of CDK1, and the other for the kinase activity assay. Immunoprecipitated CDK1 on Protein-A beads was incubated in kinase buffer with 1 mM DTT, PhosSTOP (Roche), 200 μM Mg^2+^-ATP, 3 μCi [γ-^32^P]ATP and 2 μg recombinant HI protein (Millipore) for 30 min at 30°C. The reactions were terminated by the addition of IX LDS sample buffer (NP0007, Invitrogen), and heated at 95°C for 5 min. Proteins were resolved on a 4-12% NuPAGE Bis-Tris gel (Invitrogen) and transferred to a PVDF membrane (GE Life Sciences). The blot was subjected to autoradiography (Typhoon FLA-9500) to monitor HI phosphorylation, and stained with Ponceau S to detect the amount of HI. Bands were quantified using Image J. The extent of phosphorylation of HI was quantified by first expressing the amount of radiolabeled HI as a fraction of total HI, and then normalizing this to the amount of immunoprecipitated CDK1 detected by western blotting.

### ATP binding and competition assays

The ATP binding assay was performed using the ATP affinity kit (Jena Bioscience) as per manufacturer’s protocol. Briefly, the TNN cell lysate was dialyzed overnight at 4°C against dialysis buffer (PBS, ImM EDTA, ImM DTT) with two buffer changes to remove the bound ATP-Mg^2+^. The dialyzed lysate was diluted with the binding buffer containing protease inhibitor cocktail and incubated with equilibrated control or ATP beads for 2 h at 4°C on rotor. For ATP competition, increasing amount of ATP was added in the dialyzed lysate before incubation with equilibrated control or ATP beads. The beads were washed thrice with the wash buffer for 15 minutes at 4°C on rotor and boiled in SDS gel loading buffer to elute the bound proteins.

### Recombinant 6XHis-P300 expression and purification

6XHis-tagged full-length P300 was expressed by transfection of the recombinant baculovirus into Sf21 insect ovary cells. 72 h post infection, the cells were harvested and lysed in 10 mM Tris-Cl, 500 mM NaCl, 10% glycerol, 0.1% NP-40, 15 mM imidazole, 2 mM PMSF, and 2 mM β-mercaptoethanol followed by nickel-nitriloacetic acid (Ni-NTA) affinity chromatography-based purification as described previously [59].

### Acetyltransferase Assay

In the acetyltransferase assays, 25 nM full-length P300 was incubated with 500 ng recombinant CDKl-Cyclin B1 (Merck-Millipore #14-450) in the presence or absence of 1.2 mM acetyl-CoA lithium salt (Sigma). The reaction was carried out in HAT assay buffer (50 mM Tris-HCl, pH 7.5,1 mM PMSF, 0.1 mM EDTA and 10% v/v glycerol) and 100 mM sodium butyrate at 30 °C for 90 minutes. The enzyme and acetyl-CoA were replenished twice in the reaction at an interval of 30 minutes to ensure complete acetylation of the substrate. The reaction was stopped on ice, followed by the addition of 5X SDS sample loading buffer (10 % w/v SDS, 250 mM Tris-Cl, pH 6.8, 50% glycerol, 0. 5% Bromophenol Blue, 5% β-mercaptoethanol).

### 2D-PAGE

After immunoprecipitation, the beads were solubilized in rehydration buffer (7 M urea, 2 M thiourea, 2 % CHAPS, 50 mM DTT and 2 % IPG buffer) in a final volume of 100 μl. The IPG strips (IPG ReadyStrip^™^ pH 5-8, 7 cm, Biorad #163-2004) were rehydrated overnight and IEF focusing was done with the following program:-Step 1 - 250V for 30 minutes, step 2 - 4000V for 1 h and final step 3 - 15000 Vhrs. The strips were equilibrated with 2% DTT (Sigma Aldrich) and 2.5% IAA (Sigma Aldrich) before proceeding to second dimensional gel electrophoresis on 10% SDS-PAGE gel.

### Western Blotting

Equal amount of protein was run on SDS-PAGE and transferred to PVDF membrane (Merck-Millipore). The blots were probed with the indicated primary and secondary antibodies. The chemiluminescence signal was detected using SuperSignal West Pico PLUS and SuperSignal West Femto ECL kit (Thermo Scientific) and imaged using AI600 imager (GE-Amersham).

### Microscopy

Cells were fixed using 3.7% formaldehyde for 12 minutes and permeabilized with 1% Triton. Cells were stained using 1μg/ml DAPI and 1 mg/ml Aniline Blue to visualize nucleus and cell wall respectively. Images were taken on Olympus 1X-71 Epifluorescence system with a 100X (NA 1.40) Olympus objectiveand AndoriXon EM+ EMCCD camera with using Micro manager software.

### MS/MS analysis

HEK293T cells were lysed by sonication in 20 mM HEPES containing 6 M urea and protease inhibitors (Complete, Roche). Proteins were reduced (5mM DTT), alkylated (10mM iodoacetamide), then digested with 1% trypsin (w/w) after dilution of urea to a concentration of 1M. Resulting peptides were desalted (Sep-pak C18), and acetylated peptides were enriched through immunoprecipitation with anti-Acetyl beads (PTM Scan kit, Cell Signaling Technology). Acetylated peptides were analyzed by nanoLC-MS/MS using an UltiMate 3000 RSLCnano system (Dionex, Amsterdam, The Netherlands) coupled to a LTQ-Velos Orbitrap mass spectrometer (Thermo Scientific, Bremen, Germany). Separation was performed on a C-18 column (75 μm ID x 15 cm, Reprosil C18) equilibrated in 95% solvent A (5% acetonitrile, 0.2% formic acid) and 5% solvent B (80% acetonitrile, 0.2% formic acid), using a gradient from 5 to 50% gradient of solvent B over 180 min at a flow rate of 300 nL/min. The LTQ Orbitrap Velos was operated in data-dependent acquisition mode with Xcalibur software. Survey scan MS was acquired in the Orbitrap on the 350-2,000 m/z range, with the resolution set to a value of 60,000. The 20 most intense ions survey scans were selected for fragmentation by collision-induced dissociation, and the resulting fragments were analyzed in the linear trap. Dynamic exclusion was used within 60 s to prevent repetitive selection of the same peptide. Raw mass spectrometry files were processed with the Mascot software. Peptide identification results were validated by the target-decoy approach at a FDR of 1%.

### Modelling of CDKl:Cyclin-B:ATP ternary complexes

Our computational modelling studies were initiated using a crystal structure of Cyclin-B bound CDK1 (PDB ID: 4Y72) solved by Endicott and co-workers. The 4Y72 structure was bound to an inhibitor as well as the accessory protein CKS2, both of which were removed. The resultant structure lacked ATP and C-terminal residues (resid: 290-297) for CDK1. We modelled the missing C-terminal CDK1 residues (resid: 290-297) using the loop modeling module in Modeller version 9 [60]. An ATP molecule was then introduced in the active site of the CDK1: Cyclin-B binary complex using the crystal structure of CDK2:Cyclin-A:ATP ternary complex as a reference (PDB ID: 1FIN) to create a WT CDK1:Cyclin-B:ATP ternary complex model. Specifically, we used the ATP coordinates from the 1FIN structure and inserted it at the active site of the CDKl:Cyclin-B complex retaining the same interactions of ATP with the active site aspartate-146 residue as in the 1FIN structure. After this step, we created two more mutant CDK1:Cyclin-B:ATP complex models by replacing the lysine-33 in the WT model with either glutamine (K33Q) or acetyl lysine (K33Ac). For all 3 model complexes (WT, K33Q, K33Ac), hydrogen atoms were added using the psfgen utility in VMD [61]. Then the modelled complexes were immersed in a large rectangular water box (explicit solvent model) of dimension 126×145×128 Å^3^. The net charge of the model complexes was +3 for WT and +2 for K33Q/K33Ac. Therefore 3 Clions were added to WT and 2C1— ions to K33Q/K33Ac solvent boxes to neutralize the systems.

### Equilibration protocol and Simulations

All simulations were carried out using the NAMD program [62] (version 2.9) and the CHARMM36 force field (C36 FF) [63]. Acetyl lysine was parametrized using the FFTK toolkit [64] in VMD (see parametrization procedure below). The water model used in the simulations was TIP3P. Our simulations employed periodic boundary conditions with the Particle Mesh Ewald method for describing electrostatic interactions and a switching function for van der Waals interactions (10Å switching distance and 12Å cutoff). Each of the three CDKl:Cyclin-B:ATP models (WT, K33Q and K33Ac) were subjected to an initial 10000 step energy minimization. Subsequently, the system was gradually heated at a rate 6K/lps from OK to 300K followed by an equilibration at 300K for 50 ps. The minimization and heating protocol was carried out 3 times initially with all the protein heavy atoms, ATP and Mg^2+^ fixed (with free hydrogens, solvent), then with all the protein heavy atoms other than those from modelled residues (resid 290-297), ATP and Mg^2+^ fixed, and finally with harmonic constraints of 25 kcal/mol/Å^2^ on the protein heavy atoms, ATP and Mg^2+^. These steps were then followed by constant pressure and constant temperature (NPT) equilibration runs for 150 ps to stabilize the density of the system at 1atm pressure and temperature 300 K. The pressure of the system was maintained by the Nose-Hoover method in combination with Langevin dynamics to control the temperature of the system. The NPT protocol was repeated 3 more times by lowering the harmonic constraints on the system to 12, 6 and 3 kcal/mol/Å^2^. Finally an unconstrained ∼1 ns NPT (NPT-free) run was performed at 30OK to arrive at the starting point of our production runs. The time step for all these simulations was 1 fs. In the K33Ac model, the larger size of acetyl-lysine relative to the WT lysine-33 lead to significant steric clashes during residue placement at the active site. We therefore employed three different protocols on the K33Ac model which differed in terms of constraints on ATP and E51 during the optimization, temperature equilibration, and pressure equilibration steps described above to create three distinct structurally relaxed conformations of the K33Ac ternary complex: K33Ac-1 (ATP, E51 both constrained), K33Ac-2 (ATP unconstrained, E51 constrained), and K33Ac-3 (ATP constrained, E51 unconstrained). Following the equilibration protocol, 200 ns production runs were carried out for all five systems (WT, K33Q, K33Ac-1, K33Ac-2, K33Ac-3) which were then used for subsequent analysis. Table SI summarizes all systems and simulations in the present manuscript.

### Charmm36 parameters for acetyl-lysine

In acetyl-lysine, the side-chain amino group of lysine is altered by the attachment of an acetyl (-COCH3) group at the epsilon Nitrogen (NE) position. Since no standard parameters exist in C36 FF for this modification, we parametrized the side chain acetyl head group of acetyl lysine attached to the epsilon carbon atom (CE) of Lysine (Fig S3A). This was done to give a fair description of the change in charges and bonded parameters on the side chain NZ and CE atoms, upon modifying the positively charged amino group to a neutral acetyl group. The acetylated head-group fragment is capped with an extra H atom at the lysine terminal for epsilon C to account for the absence of the bond between delta C and epsilon C. The charges on H are absorbed into the epsilon C when the acetylated head-group fragment is patched onto the lysine residue. The C36 FF comprises of non-bonded van der Waals and electrostatic interactions and covalent interactions comprising of bond, angle, and dihedrals terms. We assigned van der Waals parameters for atoms by analogy with existing C36 FF parameters. Atomic partial charges for electrostatic interactions were assigned to reproduce ab-initio interaction energies of the acetylated head-group fragment obtained from an electronic structure calculation at HF/6-31G* level of theory. In these calculations the geometry of the acetylated head-group fragment was optimized at an ab-initio HF/6-31G* level of theory. Force constants for bonded interactions (covalent bonds and angles) were obtained from Hessian calculations on the optimized geometry of the acetylated fragment. Dihedral parameters were assigned to reproduce target potential energy surfaces (PES) scans as a function of torsion angle calculated at the HF/6-31G* level of theory. Below we provide the final topology and parameters for acetyl-lysine residue derived using FFTK.

CHARMM36 Topology file

RESIALY 0.00 GROUP !

ATOM N NH1 −0.47 ! |

ATOM HN H 0.31 ! HN-N

ATOM CA CT1 0.07 ! | HB1 HG1 HD1 HE1 HZ OCD HM1

ATOM HA HB1 0.09 ! | | | | | | | |

GROUP ! HA-CA-CB-CG-CD-CE-NZ-CO-CM-HM3

ATOM CB CT2 −0.18 ! | | | | | |

ATOM HB1 HA2 0.09! | HB2HG2HD2HE2 HM2

ATOM HB2 HA2 0.09 ! 0=C

GROUP ! |

ATOM CG CT2 −0.18 !

ATOM HG1 HA2 0.09

ATOM HG2 HA2 0.09

GROUP

ATOM CD CT2 −0.18

ATOM HD1 HA2 0.09

ATOM HD2 HA2 0.09

GROUP

ATOM CE CT2 −0.598

ATOM HE1 HA2 0.227

ATOM HE2 HA2 0.227

GROUP

ATOM NZ NZ −0.244

ATOM HZ HZ 0.247

GROUP

ATOM CM CM −0.418

ATOM HM1 HM 0.133

ATOM HM2 HM 0.133

ATOM HM3 HM 0.133

GROUP

ATOM CO CO 0.822

ATOM OCD OCD −0.662

GROUP

ATOM C C 0.51

ATOMO O −0.51

BOND CB CA CGCB CD CG CE CD NZ CE CE HE1 CE HE2

BOND NHN NCA C CA

BOND C +N CAHA CB HB1 CB HB2 CG HG1

BOND CG HG2 CD HD1 CD HD2

DOUBLE 0 C

BOND NZ HZ

BOND CO NZ CM CO CM HM1 CM HM2 CM HM3

DOUBLE OCD CO

IMPRN-CCAHN CCA+NO

CMAP -C N CA C N CA C +N

DONOR HNN

DONOR HZ

ACCEPTOR 0 C

ACCEPTOR OCD NZ

CHARMM36 Parameter file:

BONDS

!V(bond) = Kb(b - b0)**2

!

!Kb: kcal/mole/A**2

!b0: A

!

!atom type Kb bO

!

NZ HZ 489.677 1.010

NZ CO 462.890 1.361

NZ CT2 417.386 1.418

CO OCD 767.853 1.236

CO CM 296.309 1.536

CE HE1 345.280 1.090

CE HE2 345.280 1.090

CM HM 363.627 1.093

ANGLES

!

!V(angle) = Ktheta(Theta - Theta0)**2

!

!V(Urey-Bradley) = Kub(S - S0)**2

!

IKtheta: kcal/mole/rad**2

!ThetaO: degrees

!Kub: kcal/mole/A**2 (Urey-Bradley)

ISO: A i

!atom types Ktheta ThetaO Kub SO

!

!

NZ CT2 HA2 71.626 107.750

NZ CO CM 117.947 115.101

NZ CO OCD 122.322 122.967

HZ NZ CT2 59.174 119.740

HZ NZ CO 75.065 124.254

CO CM HM 66.097 110.245

CO NZ CT2 96.613 117.960

HE1 CT2 HE2 78.469 109.472

CM CO OCD 113.730 124.779

HM CM HM 45.989 107.983

CT2 CT2 NZ 70.000 113.5000 ! ALLOW ALI PEP POL ARO! from NH1 CT1

CT2, for lactams, adm jr.

DIHEDRALS

!

!V(dihedral) = Kchi(1 + cos(n(chi) - delta))

!

!Kchi: kcal/mole

!n: multiplicity Idelta:

!degrees

!

!atom types Kchi n delta

!

HZ NZ CO CM 1.2250 2 0.00

HZ NZ CO CM 1.2100 3 0.00

CT2 NZ CO OCD 0.5240 3 0.00

CT2 NZ CO OCD 2.6160 2 180.00

OCD CO CM HM 2.9810 2 0.00

OCD CO CM HM 0.3220 3 180.00

HZ NZ CO OCD 0.7860 3 180.00

HZ NZ CO OCD 2.3940 2 180.00

CT2 NZ CO CM 2.4350 3 180.00

CT2 NZ CO CM 3.0000 2 180.00

NZ CO CM HM 0.4860 3 180.00

NZ CO CM HM 2.4860 2 0.00

HZ NZ CT2 HA2 1.7510 2 0.00

HZ NZ CT2 HA2 0.0500 3 0.00

CO NZ CT2 HA2 0.0450 2 0.00

CO NZ CT2 HA2 0.0160 3 0.00

CO NZ CT2 HA1 0.0450 2 0.00

CT2 CT2 NZ HZ 0.0000 1 0.00 ! ALLOW PEP ! from H NH1 CT2 CT3, for lactams, adm jr.

CT2 CT2 NZ CO 1.8000 1 0.00 ! ALLOW PEP ! from CT2 CT1 NH1 C, for lactams, adm jr.

IMPROPER

!

!V(improper) = Kpsi(psi - psi0)**2

!

!Kpsi: kcal/mole/rad**2

!IpsiO: degrees

!note that the second column of numbers (0) is ignored

!

!atom types Kpsi psi0

!

NONBONDED nbxmod 5 atom cdiel shift vatom vdistance vswitch –

cutnb 14.0 ctofnb 12.0 ctonnb 10.0 eps 1.0 el4fac 1.0 wmin 1.5

!

!V(Lennard-Jones) = Eps,i,j[(Rmin,i,j/ri,j)**12 - 2(Rmin,i,j/ri,j)**6]

!

!epsilon: kcal/mole, Eps,i,j = sqrt(eps,i * eps,j)

!Rmin/2: A, Rmin,i,j = Rmin/2,i + Rmin/2,j

!

!atom ignored epsilon Rmin/2 ignored eps,1-4 Rmin/2,1-4

!

CE 0.0 −0.056000 2.010000 0.0 −0.010000 1.900000

CM 0.0 −0.078000 2.040000 0.0 −0.010000 1.900000

CO 0.0 −0.110000 2.000000

HE 0.0 −0.034000 1.340000

HM 0.0 −0.024000 1.340000

HZ 0.0 −0.046000 0.224500

NZ 0.0 −0.200000 1.850000 0.0 −0.200000 1.550000

OCD 0.0 −0.120000 1.700000 0.0 −0.120000 1.400000

END

### Statistical analysis

Statistical analyses for biochemical data were performed using GraphPad Prism (Version 6.0) and Microsoft Excel 2013. For comparing two groups, Student’s t-test was performed. Statistical analyses for CMD data were performed using MATLAB (version R2014a).

## Acknowledgements

The following laboratories were supported by the indicated research grants. UK (TIFR/DAE 12P0122 and CEFIPRA 4503-1), RV (TIFR/DAE 12P0155), MM (TIFR/DAE 12P0128), RB (HFSP RGP0025/2016), TKK (JNCASR and Sir JC Bose Fellowship, DST-India; SR/S2/JCB-28/2010), AGP (Investissement d’Avenir Infrastructures Nationales en Biologie et Sant program; ProFI, Proteomics French Infrastructure project, ANR-10-INBS-08). We thank Prof. Matthias Wymann (University of Basel, Switzerland) for providing a-FSBA antibodies.

## Author Contributions

UKS/RV conceptualized the project with inputs from SD, MM, TKK, RB & AGP. SD, SR, NK, MB, AF, RS, SG, CB, SK, KKV, AGP, RV and UKS performed the experiments. SD, RV and UKS wrote the manuscript All authors have contributed towards analysis and interpretation of data.

## Supplemental figure legends

**Figure SI - CDK1 is acetylated at lysine-33 in-vivo.** (A) Flow chart of the protocol for affinity purification of a-CDKl-K33Ac antibody. (B) Sequence of the peptide used for generating α-CDKl-K33Ac antibody and the high conservation of that region across species. (C) ELISA of purified antibodies after the 3^rd^ immunization on acetylated and non-acetylated peptides shows high specificity of α-CDKl-K33Ac antibodies. (D) Probing for acetylation of HA immunoprecipitates from cells expressing CDK1-WT-HA or -K33R-HA shows specificity of α-CDKl-K33Ac antibody. (E) 2D-PAGE of immunoprecipitated endogenous CDK1 from mouse embryonic fibroblasts (MEFs) and immunoblot with a-CDKl-K33Ac antibodies confirms acetylation. (F) In-vitro acetylation assay using recombinant CDK1:Cyclin-Bl complex and 6XHis-P300 shows P300 dependent acetylation of CDK1. (G) Immunoblot of HA immunoprecipitates from cells co-transfected with SIRT1 shows CDK1-SIRT1 interaction.

**Figure S2 - CDK1 lysine-33 acetylation abolishes kinase activity but does not affect cell cycle progression.** (A) HI phosphorylation by CDK1-WT, -K33Q and -K33R shows lysine-33 mutation abrogates kinase activity. (B) Flow cytometry analysis of cells used for the kinase assay in Fig 2A and Fig 2B. (C) Flow cytometry analysis of HeLa-pTetON cells transfected with CDK1-WT or -K33Q shows no change in cell cycle profile. (D) Immunoblot showing the expression of CDK1-WT and -K33Q for cells used for flow cytometry analysis in (C).

**Figure S3 - CMD simulations for WT, K33Q and K33Ac-1,-2,-3 CDKl:Cyclin-B:ATP ternary complexes and the impact of CDK1 lysine-33 mutation/acetylation on ATP-active site and ATP-PSTAIRE helix interactions.** (A) Acetylated head-group fragment for which Charmm36 parameters were derived using FFTK. (B-K) Modeled pre-simulation structures for WT, K33Q and K33Ac-1,-2,-3 CDKl:Cyclin-B:ATP ternary complexes obtained from the crystal structure using the workflow depicted in Fig 3A. (B-F) show CDK1 (multi-color) and Cyclin-B (orange) in secondary structure representation. ATP bound at the CDK1 active site is shown in atomic van der Waals spheres representation. (G-K) View of the active site showing key catalytic triad residues, lysine-33 (K33)/glutamine-33 (Q33)/acetyl-lysine (K33Ac), glutamate-51 (E51) and aspartate-146 (D146) interacting with the bound ATP and Mg^2+^ (green). (L) Non-bonded interaction energies of ATP with acidic (left panel) and basic (right panel) residues within 3-7A in WT, K33Q and K33Ac CDK1:Cyclin-B:ATP systems are favorable. Error bars represent SD. (M-0) Interactions of K33 with ATP and E51 during 200 ns MD trajectories for the WT CDKl:Cyclin-B:ATP complex (M) Non-bonded (electrostatic + van der Waals) interaction energies, (N) Distance between the sidechain amino nitrogen of lysine and the two carboxylate oxygens of glutamate, (0) distance between the sidechain amino nitrogen of lysine and three of the closest oxygens of the ATP phosphate tail. (P-R) Interactions between the PSTAIRE helix and ATP do not change in WT, K33Q and K33Ac CDKl:Cyclin-B:ATP systems. (P) Distance between the PSTAIRE helix (COM) and ATP (COM), (Q) distance between E51 (COM) and ATP (COM), (R) Non-bonded interaction energy between ATP and E51.

**Figure S4 - PSTAIRE helix and T-loop undergo a conformational change upon CDK1 lysine-33 mutation/acetylation.** (A) Relative orientation of PSTAIRE helix in a representative snapshot of CDK1-K33Q complex relative to its orientation in the first frame for CDK1-WT. (B) Rotation of PSTAIRE helix during 200 ns MD trajectories for WT, K33Q and K33Ac CDKl:Cyclin-B:ATP systems. The angle made by the PSTAIRE helix axis vector during each frame relative to its direction in the first frame of the CDK1-WT trajectory (CDK1 backbone alignment) was computed. (C) Percentage solvent accessibility (fractional SASA of T-loop with and without surrounding protein) computed for the T-loop segment (residues 152-171) along the 200 ns MD trajectory. (D) Representative conformations of the T-loop during 200 ns MD production runs for WT, K33Q and K33Ac CDKl:Cyclin-B:ATP systems. The full CDKl:Cyclin-B complex is shown on the left and the view on the right zooms in on the T-loop and its neighbourhood.

**Figure S5 - CDK1 acetylation at lysine-33 does not affect ATP binding but reduces Cyclin-Bl interaction.** (A) Probing for pull-down of CDK1-WT-HA using blank or different ATP conjugated agarose beads shows highest binding to 6-AH beads. (B) Flow cytometry analysis of cells used for the ATP binding assay in Fig 5B and Fig 5C. (C) Probing for CDK1-WT/-K33Q/-K33R-HA binding (pull-down) by 6-AH beads shows no change in steady-state binding. S6K binding was used for normalization. (D, E) ATP competition assay of CDK1-WT/-K33Q/-K33R-HA using blank or 6-AH beads shows small differences. Coomassie staining was used for normalization. Error bars represent SEM for three experimental replicates. (F, G) Reversible co-immunoprecitations of CDK1-WT/-K33Q/-K33R/-D146N-HA and Myc-Cyclin-Bl shows reduced CDKl:Cyclin-Bl binding upon K33Q mutation. Immunoprecipitation of complexes with (F) α-Myc and (G) α-HA antibodies. * marks the IgG heavy chain. (H) Flow cytometry analysis of cells used for assaying CDK1 acetylation in Fig 5H.

**Figure S6 - Loss of CDK1 lysine-33 charge state affects surface interactions with Cyclin-B.** (A, B) Histograms of interfacial salt bridges between (A) sidechains and (B) side chain-backbone of acidic and basic residues of CDK1 and Cyclin-B show reduced number of salt-bridges in K33Q and K33Ac CDK1:Cyclin-B:ATP systems relative to WT.

**Figure S7 - Phenotypic characterization of fission yeast *cdc2-33* and *cdc2-L7* mutants in presence of plasmid expressing Cdc2-WT/-K33Q/-K33R.** (A) Quantification of cell length measurements observed in *cdc2-33* and *cdc2-L7* strains transformed with indicated plasmid at permissive temperature of 25°C. (B) Quantification of binucleate cells after 16 h of induction at 25°C. (C) Quantification of binucleate cells observed when shifted to restrictive temperature of 36°C for 4 h after 16 h of induction. Error bar indicates SEM for three technical replicates of independent experiments where n=500 for each sample. (D) Representative images of DAPI and aniline blue co-stained cells at 25°C (top panel) and 36°C (lower panel) respectively. (E) Plate viability assay of *cdc2-L7* strains over-expressing indicated plasmids at 25°C and 36°C respectively.

## Supplementary tables

**Table SI** - Summary of all CDK:Cyclin-B:ATP ternary complexes and their simulation trajectories.

**Table S2** - Salt bridges between acidic and basic residues of CDK1 and Cyclin-B within 4 Å of the binding interface formed during the first 1 ns of the 200 ns MD production (NVT) simulations for the 5 different model complexes. Salt bridges in the mutant (K33Q and K33Ac models) complexes, also present in the WT, are highlighted in green while those that are distinct from WT are highlighted in red.

